# HOW CONCERNING IS A SARS-COV-2 VARIANT OF CONCERN? COMPUTATIONAL PREDICTIONS AND THE VARIANTS LABELING SYSTEM

**DOI:** 10.1101/2022.01.31.478425

**Authors:** Dana Ashoor, Maryam Marzouq, Khaled Trabelsi, Sadok Chlif, Nasser Abotalib, Noureddine Ben Khalaf, Ahmed R. Ramadan, M-Dahmani Fathallah

## Abstract

We herein report a study to evaluate the use of computational prediction of SARS-CoV-2 genetic variations in improving the current variants labeling system. First, we reviewed the basis of the system developed by the World Health Organization (WHO) for the labeling of SARS-CoV-2 genetic variants and the adaptations made to it by the United States Center of Diseases Control (CDC). We observed that the labeling system is based upon the virus’ major attributes. However, we found that the labeling criteria of the SARS-CoV-2 variants derived from these attributes are not accurately defined and are used differently by the two health management agencies. Consequently, discrepancies exist between the labels given by WHO and CDC to same variants. Our observations suggest that giving the VOC label to a new variant is premature and might not be appropriate. Therefore, we carried out a comparative computational study to predict the effects of the mutations on the virus structure and functions of five VOCs. By linking these data to the criteria used by WHO and the CDC for variant labeling, we ascertained that comparative computational predictions of the impact of genetic variations are a better ground for rapid and more accurate labelling of SARS-CoV-2 variants. We propose to label all emergent variants VUM or VBM and to carry out computational predictive studies and thorough variants comparison, upon which more appropriate and informative labels can be attributed. Furthermore, harmonization of the variants labeling system would be globally beneficial to communicate about and fight COVID-19 pandemic.

## 1. INTRODUCTION

The new coronavirus SARS-CoV-2 that emerged late 2019 is still causing a pandemic of Severe Acute Respiratory Syndrome or COVID-19. The pandemic affected over 370 million people and claimed more than 5.5 million (Johns-Hopkins-University, 2022), thus posing a difficult challenge to the scientific and health care communities throughout the world (Malik et al., 2020; Tutelyan et al., 2020; World-Economic-Forum, 2020). Indeed, SARS-CoV-2 is a positive RNA virus that is constantly evolving through the accumulation of various type of mutations (Zhao et al., 2020; Banoun, 2021; Majumdar and Niyogi, 2021; Singh et al., 2021). Even though the majority of these mutations do not affect the virus infectious properties and have no real impact on the progress of the pandemic (Ashoor et al., 2021), some may enhanced specific viral attributes that give the virus a selective advantage (Gobeil et al., 2021). Any new variant endowed with selective advantage(s) would favor the virus persistence and nurture the pandemic. Therefore, watching and predicting how the pandemic evolves and communicate it to the public is of paramount importance. The global surveillance of the pandemic is based on multidisciplinary approaches including epidemiological, genetic, structural, and clinical data (Agency, 2021; Campbell et al., 2021; England, 2021; France, 2021). This involves the use of a set of relevant criteria to categorize the variants. Toward this-end, all international and national health and sanitary authorities have set various strategies to control the evolution of the SARS-CoV-2 pandemic. In May 31, 2021, the World Health Organization (WHO) announced a labeling system to categorize the variants into different levels of priority to better organize the global monitoring and research, and ultimately organize the “infodemic” and communicate more effectively with the public about the adequate response to the emergence of new variants of SARS-CoV-2 (https://www.who.int/). WHO has first developed a system to facilitate naming SARS-CoV-2 variants in addition to the existing nomenclature systems for naming and tracking SARS-CoV-2 genetic lineages established by GISAID (https://www.gisaid.org/), Nextstrain (https://nextstrain.org/) and Pango (https://cov-lineages.org/).These nomenclatures are mostly used by the scientific research community. For practical reasons (particularly to ease the communication), the WHO has settled to name the SARS-CoV-2 emerging variants using the Greek letter alphabet sequence (α, ß, γ, δ.….). Since the SARS-CoV-2 virus is showing high genetic variability (Toyoshima et al., 2020; Yazdani et al., 2021; Dubey et al., 2022), WHO has established a labeling system for the variants into variant of concern (VOC), variants of interest (VOI) and variant under monitoring (VUM). This labeling system is based upon definitions related to variant phenotypic attributes such as transmissibility, disease presentation, effect on current diagnostic tests and response to available vaccines. This system was set to prompt and harmonize the actions needed to control the spread of a given variant. While the variants that emerged sequentially were named α, β, γ and δ, the last emerging one was named *omicron* which does not comply with the sequence of the Greek alphabetic order that call for naming this variant “*Nu*”. WHO deemed that “*Nu*” is prone to confusion with “New”. *Xi* was also not used because it is a common surname in Asia. Therefore, this variant was named “*Omicron*”. Of interest is that the American Center for disease control has also adopted this system of variant classification and labelling but added more labels and labeling criteria and a different labels’ change policy (https://www.cdc.gov/). Indeed, while keeping the labels, variant of concern (VOC), variant of interest (VOI) and variant under monitoring (VUM) (calling the later “VBM” for Variant Being Monitored). In addition, the CDC uses an extra label that is the “Variant Of High Consequences” (VOHC). As a result, some of the currently known variants are given different labels by each agency. Indeed, according to the CDC there is no SARS-CoV-2 variant labeled VOI as of December 1, 2021. Furthermore, variants α, β and γ that are currently labelled VOC by WHO, have been deescalated to the VBM label by the CDC as of September 2021 (https://www.cdc.gov/). These discrepancies reflect different views on the labeling of SARS-CoV-2 variants and consequently the use of the labels to set public health actions. On another hand, in both systems, variants labels can change with more data accumulating for a particular variant.

In this work, we have undertaken an evaluation of the system developed by WHO and adapted by the CDC to label the SARS-CoV-2 variants. We carried out a review of the classifications criteria and analyzed how WHO and the CDC use these criteria to label the SARS-CoV-2 variants. Then we carried out an exhaustive comparative computational study of the S protein mutations that characterize the five SARS-CoV-2 VOCs. We concluded that computational predictions provide a good ground of evidences for a rapid and more accurate labeling system.

## 2. MATERIALS & METHODS

### 2.1. Data mining and information sources

We retrieved the genetic, epidemiological and clinical data on the variants available as of December 15, 2021 from primary and secondary sources including the GISAID (https://www.gisaid.org/) and the Variants (http://variants.org) data banks. We collected the information on variants naming and labelling from the following sources: WHO: https://www.who.int/en/activities/tracking-SARS-CoV-2-variants/. CDC: https://www.cdc.gov/coronavirus/2019-ncov/variants/variant-info.html#anchor_1632154493691. and PANGO Lineages.

### 2.2. Variant sequences retrieval and solvent exposure analysis

SARS-CoV-2 spike protein extracellular domain amino acid sequence was obtained from the National Center for Biological Information (NCBI) protein ID: YP_009724390.1. Variants-specific mutations were introduced to the collected sequence based on the list published on (https://covariants.org/). The sequences corresponding to the different variants (*alpha*, *beta*, *gamma*, *delta* and *omicron*) were analyzed for solvent exposure and possible epitope residues using Sequential B-Cell Epitope Predictor server (BepiPred-2.0 server: https://services.healthtech.dtu.dk/service.php?BepiPred-2.0). BepiPred-2.0 is based on a random forest algorithm trained on epitopes annotated from antibody-antigen protein structures (Jespersen et al., 2017).

### 2.3. SARS CoV-2 spike protein furin cut site (FCS) loop modeling

For the loop modeling, Phyre2 web server (http://www.sbg.bio.ic.ac.uk/phyre2/html/page.cgi?id=index). (Kelley et al., 2015) was used to generate *alpha, beta, gamma, delta* and *omicron* variants 3D models of the extracellular spike monomers, the results were saved and visualized on PyMOL (DeLano, 2002).

### 2.4. Model quality assessment

As a quality assessment for the generated models, crystalized model of the spike protein (PDB ID 6VXX 2.80A° version 2.4) was downloaded from the RCSB database (https://www.rcsb.org). The structure was cleaned of water and heteroatoms, the complex was split and a PBD file for a monomer chain was created (chain B) and saved using PyMOl software. This monomer chain B was used as reference model for the sequence generated models. To define the common contact map between the crystal structures and the generated models, CMView (Vehlow et al., 2011) with the following parameters (contact type: Ca; Distance cut-off: 8.0; Needleman-Wunsch alignment), was used. Different contact maps were established between the crystal structure and the models and the common contact percentage was calculated. Higher common contacts indicate more structural similarity and hence the models are suitable for further analysis. In addition, Tm align (https://zhanggroup.org/TM-align/) (Zhang and Skolnick, 2005) was used to calculate TM-score value for each model. The superimposition Root Mean Square Deviation (RMSD) was calculated using PyMOL. Low RMSD and TM-scoring between 0.5 and 1.0 indicate that the two compared structures (the crystal and the model) has about the same fold.

### 2.5. Mutational analysis: The effect of mutation on the interaction with ACE2 receptor

To analyze the effect of different RBD mutations on different SARS-CoV-2 variants interaction with ACE2 receptor, PDB ID 6LZG structure was used as a model. The effect of single and accumulated mutations were evaluated by calculating changes in binding affinity (ΔΔG) upon single or multiple mutations using MutaBind2 server (https://lilab.jysw.suda.edu.cn/research/mutabind2/) (Zhang et al., 2020). In addition, the server provide a structural model that was used to analyze polar interactions by LigPlot+ software (Laskowski and Swindells, 2011).

## 3. RESULTS

### 3.1. Study of SARS-CoV-2 variants labeling system

Expert groups at the World Health Organization and the American Center of Disease Control have developed similar labeling systems to classify the new emergent SARS-CoV-2 variants. We retrieved a total of 24 different criteria from the working definitions elaborated by WHO and the CDC to give a particular label to a new variant. These criteria are derived from six viral attributes (Table 1). To each attribute, correspond a set of criteria that are formulated differently by each agency (see Table 1) who considers different criteria in their working definition of each label. Figure 1 shows that the two agencies use common (VOC, VOI) and different (VUM, VBM and VOHC) labels. Each agency uses generally different combination of criteria to give a variant a specific label except in four instances shown in Figure 1 where they use overlapping criteria for the same label (VOC and VOI). Consequently, there are discrepancies in the labels of currently active variants. Indeed, while WHO is currently labeling variants α, ß,γ, δ and *omicron* as VOC, for the CDC, only variant *omicron* is labeled VOC. Indeed, the CDC declassified variants α, β, γ, and δ to VBM. In addition, there are no variants currently labelled VOI by the CDC. Figure 1 illustrates the differential use by WHO and the CDC of the viral attributes-derived criteria to label the SARS-CoV-2 genetic variants.

**Figure 1:**
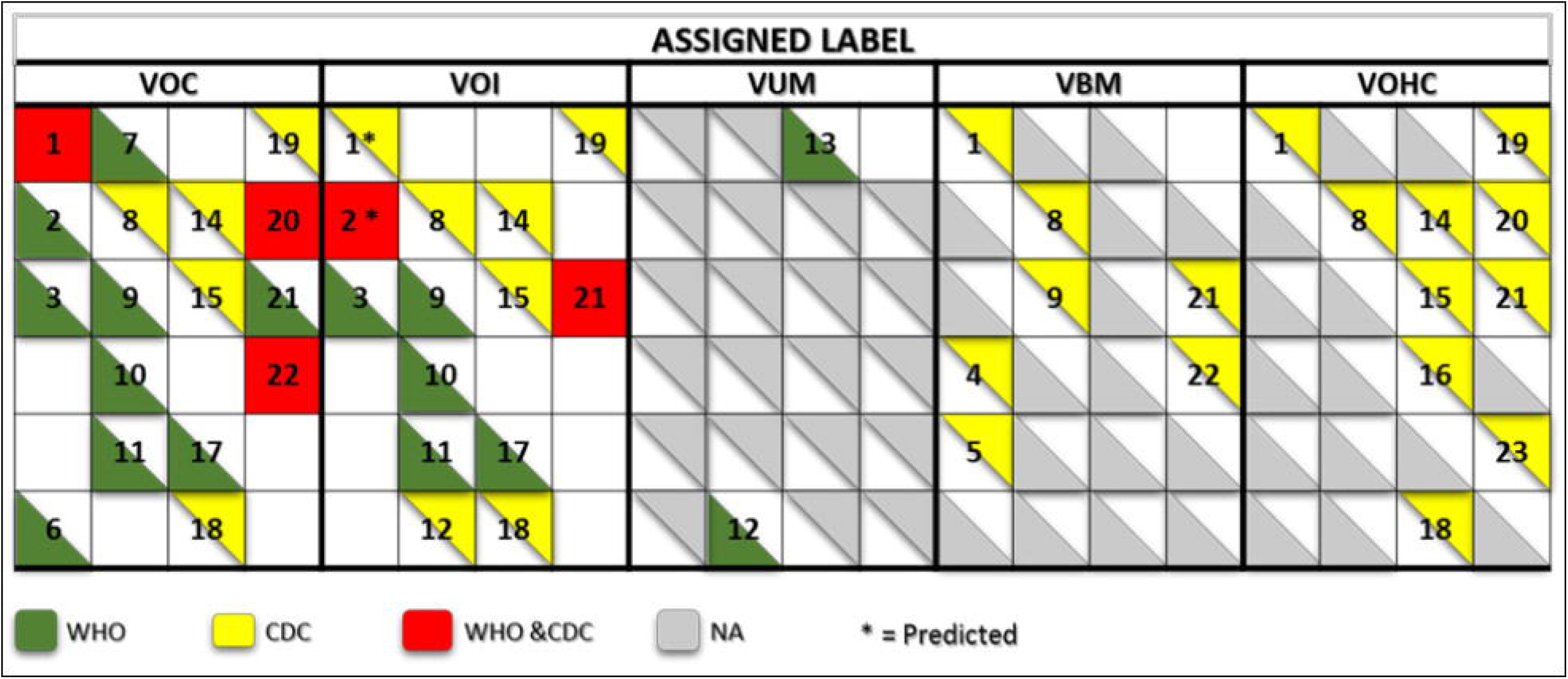
Matrix showing the commonalities and discrepancies of the criteria used by WHO and the CDC to label SARS CoV-2 variants. The colors green and yellow indicate the criteria used by WHO and the CDC respectively. The red indicates the criteria used by both agencies for a same label. The numbers correspond to the labeling criteria displayed in Table 1.

**Table 1:** SARS-CoV-2 variants attributes and derived criteria used by the World Health organization (WHO) and the American Center of Disease Control (CDC) for the labeling of emergent variants.

| Elements of SARS-CoV-2 Variants Labeling Systems |  | Labeling criteria |  |
| --- | --- | --- | --- |
| Viral attributes |  |  |  |
| Transmissibility |  | 1 | Increase in transmissibility |
|  |  | 2 | Changes in transmissibility |
|  |  | 3 | Significant community transmission or multiple COVID-19 clusters |
|  |  | 4 | Variants circulating in a limited geographic area |
|  |  | 5 | Very low circulation level or no longer detected/Limited prevalence |
| Disease presentation |  | 6 | Increase in virulence |
|  |  | 7 | Change in clinical presentation |
|  |  | 8 | Increase in disease severity |
|  |  | 9 | Changes in disease severity |
| Epidemiological impact |  | 10 | Increasing number of cases over time/prevalence |
|  |  | 11 | Other apparent epidemiological impacts |
|  |  | 12 | Genetic changes that are suspected to affect virus characteristics |
|  |  | 13 | Unclear evidence of phenotypic or epidemiological impact |
| Impact on therapy and Diagnostic |  | 14 | Reduced effectiveness of treatments/ class of therapies |
|  |  | 15 | Impact diagnostic tests |
|  |  | 16 | Diagnostic detection failures |
|  |  | 17 | Diagnostic or therapeutic escape |
| Impact on immunity & vaccination |  | 18 | significant reduction of previous or post vaccine immunity/Reduced Neutralization by antibodies |
|  |  | 19 | Reduced Neutralization by antibodies |
|  |  | 20 | Decreased vaccines effectiveness |
|  |  | 21 | Immune escape |
| Impact on public health measure |  | 22 | Decrease in effectiveness of public health and social measures |
|  |  | 23 | Reduced previously authorized medical countermeasures |
|  |  | 24 | Imminence of risk to public health |

### 3.2. Attributes of the SARS-CoV-2 major Variants Of Concern

#### 3.2.1. Epidemiological impact and transmissibility

While epidemiological data are available for variants α, β, γ and δ, data on the *omicron* variant are limited and incomplete. This is mainly because not enough time has elapsed since the emergence of this variant to allow enough data accumulation and meaningful analysis. Table 2 summarizes the initial pieces of data available as of December 25, 2021 on the *omicron* variant. For this variant, the early estimation of transmissibility increase is in the range of 3 to six. However, the previous data observed with the transmissibility of variant delta that became dominant worldwide and the currently observed high spreading /incidence of the *omicron* variant makes the prediction of its transmissibility a 100-fold higher than the *delta* variant plausible (Rao and Singh, 2021).

**Table 2:**
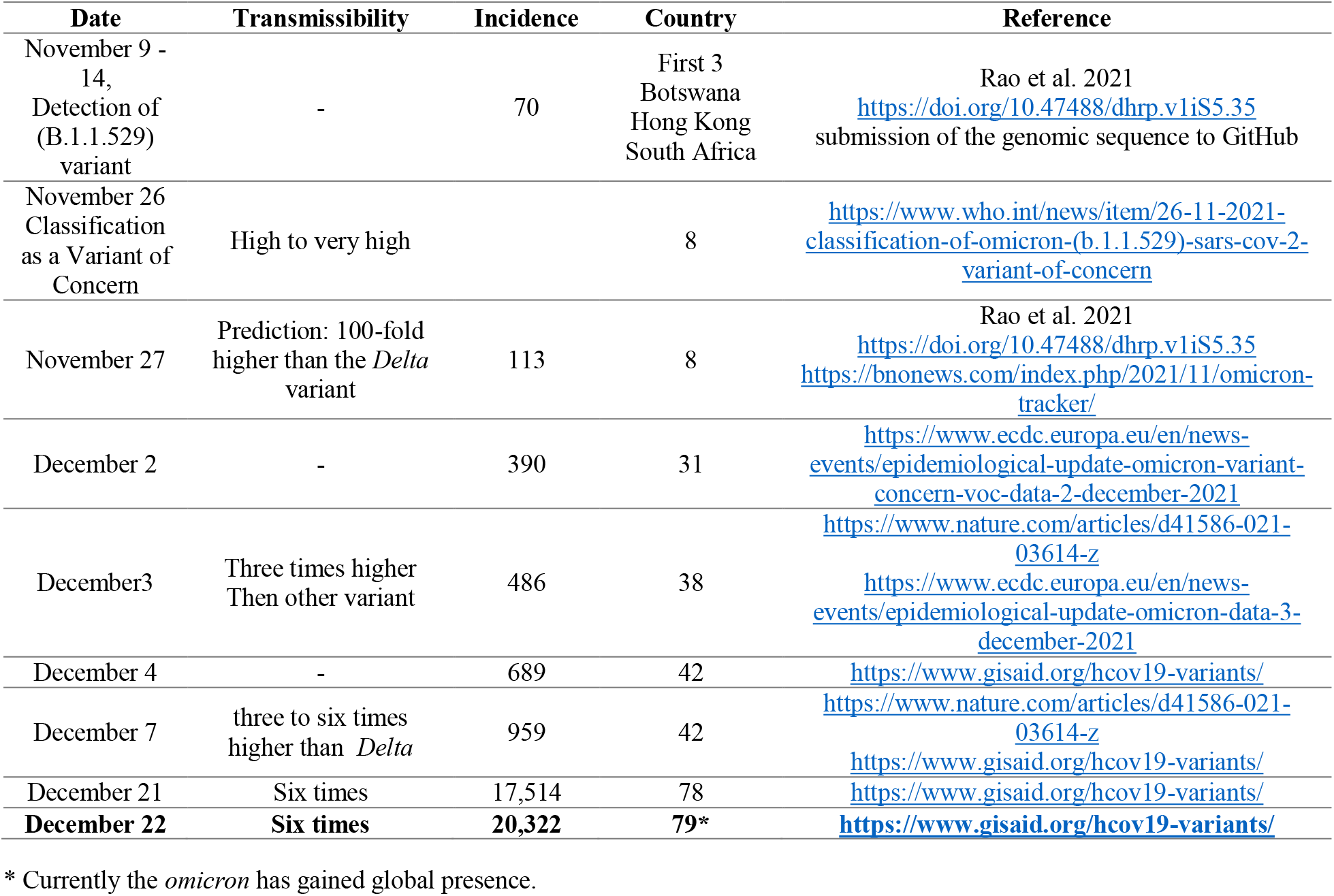
Global Spreading of SARS-CoV-2 *omicron* variant (B.1.1.529)

#### 3.2.2. Disease presentation and impact on therapy and diagnostic

COVID-19 has three clinical presentation forms mild, moderate and severe. WHO and the CDC use four labeling criteria related to clinical presentation and four others pertaining to the impact a given variant on therapy and diagnostic. The two agencies use different formulation for all of these criteria (see Table 1) and use different combination of these criteria to label SARS-CoV-2 variants (see Figure1). There is no explicit mention to the disease forms in the variant labeling criteria related to disease presentation in both WHO and CDC variant-labeling usage.

#### 3.2.3. Impact on immunity, vaccination and public health measures

WHO and the CDC use also different formulation for the criteria related to these attributes and use them differently to attribute of a specific label (Table 1, Figue1). For instance, to WHO immune escape is a criterion used to label a variant VOI and VOC. However, the CDC uses it for labeling a variant VOI, VBM and VOHC. *Omicron* has been shown to have extensive but incomplete escape of Pfizer BNT162b2 vaccine (Cele et al., 2021) thus fulfilling the criterion of decreased vaccine effectiveness. For the criterion related to the evaluation of an imminent risk to public health a variant can pose, only the CDC uses it for the VOHC label. Table 3 show how the five major SARS-CoV-2 variants fulfill the criteria used by WHO and the CDC for variants labeling regardless of the actual labels given by each agency.

**Table 3:**
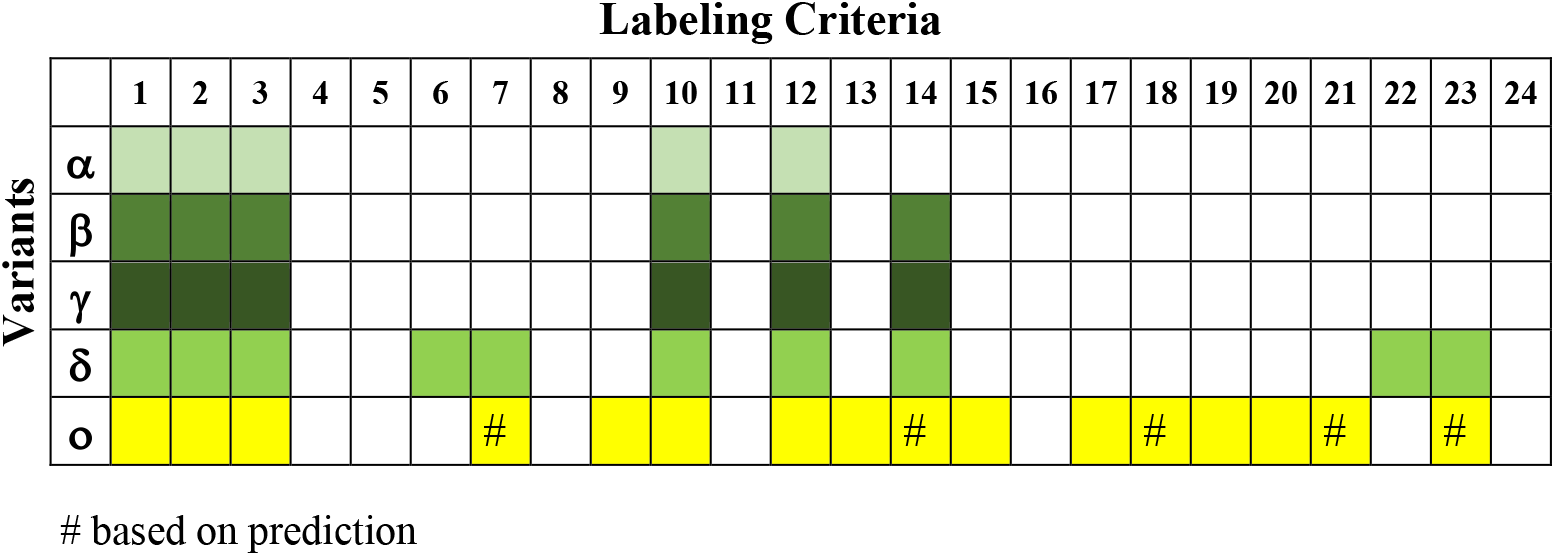
Actual application of the criteria formulated by WHO and the CDC to the five major SARS-CoV-2 variants.

### 3.3. Genetic variations data

Figure 2 shows a comparative mapping of the mutation profile of the *omicron* variant with those of the *alpha, beta, gamma* and *delta* variants. Most of the mutations affect the SARS-CoV-2 S protein. *omicron* display 63 different mutations as compared to the Wuhan strain. Thirty-six mutations occurred in the S spike protein and fifteen are clustered in the RBD region. The representation of the mutations in the S protein 3D models of SARS-CoV-2 five VOCs shows that most of the mutations map to the solvent exposed regions (Figure 3 and Supplementary Table 1).

**Figure 2.**
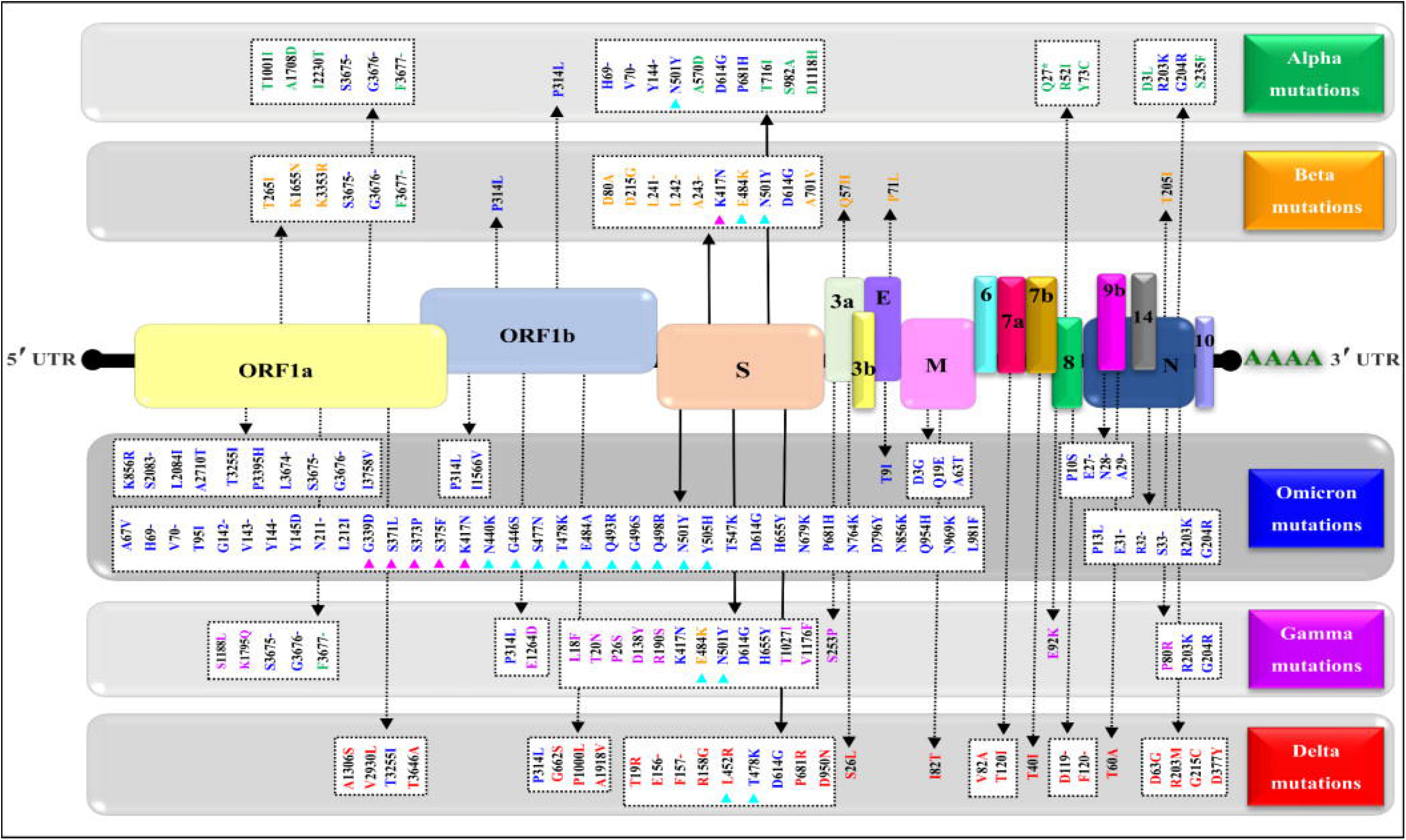
Mapping of the nonsynonymous mutations’ characteristics of the SARS-CoV-2 five variants. A typical genomic organization of SARS-CoV-2 contains the following: 5’ end UTR; Open reading frames: ORF 1a and ORF 1b; the structural genes coding for the Spike (S) protein, the Envelope (E), the Membrane (M), and the Nucleocapsid. The accessory genes such as (3a, 3b, 6, 7a, 7b, 8, 10 &14) are distributed among the structural genes. The 3’ end UTR and follows the poly (A) tail. The green, yellow, blue purple and red show respectively the synonymous mutations characteristic of variant the *Alpha* (α) variant (B.1.1.7), *Beta* (β) variant (B.1.351), *Omicron* (o) variant (B.1.1.529), *Gamma* (γ) Variant (P.1) and *Delta* (δ) variant (B.1.617.2). The (-) represents the deletion, (*) represents the stop codon, magenta triangles indicate variations in the receptor-binding domain (RBD) and cyan triangles indicate variations in the receptor-binding motif (RBM).

**Figure 3.**
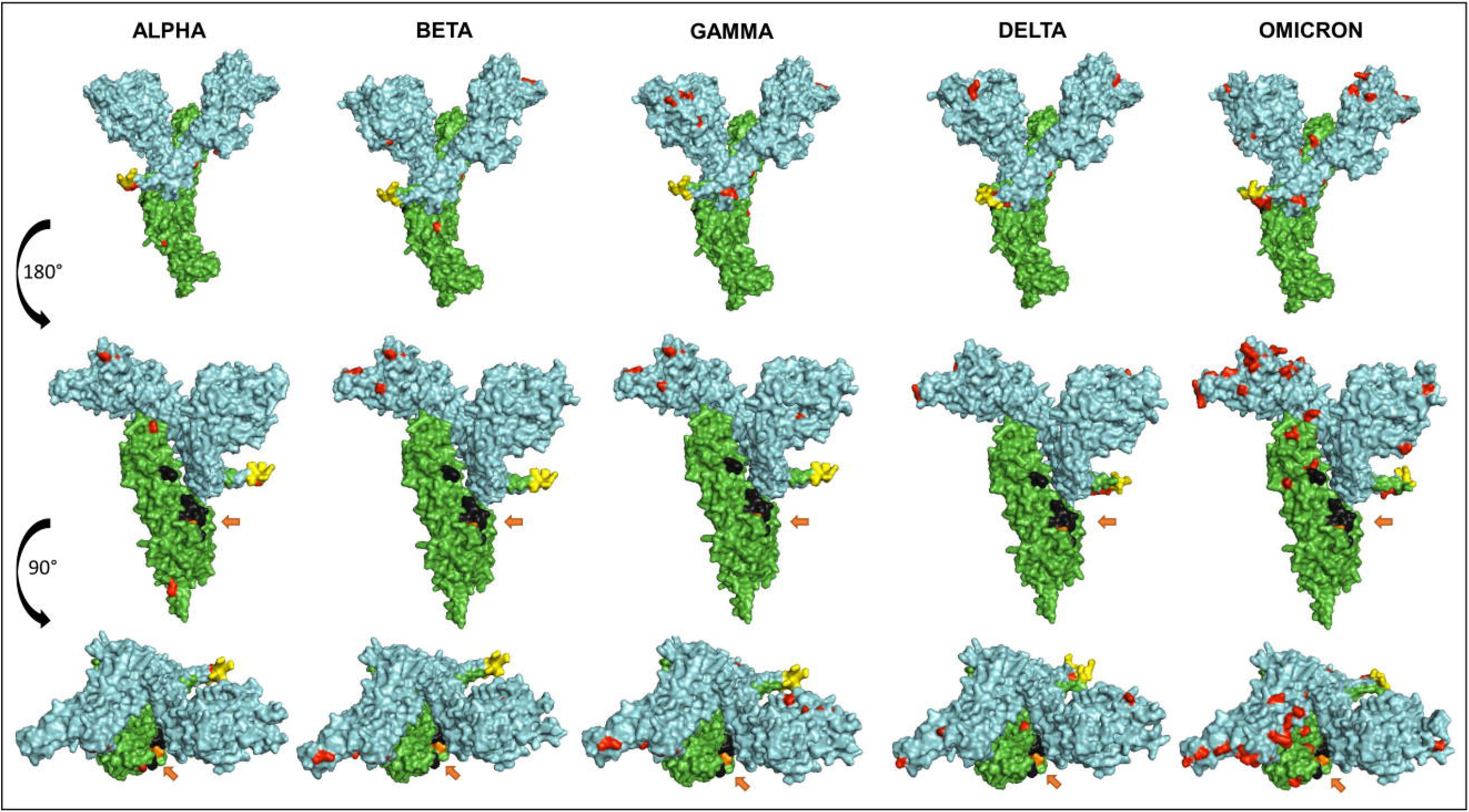
Representation of the surface models of SARS CoV-2 variant S spike protein (monomer). Side views (upper and middle rows), Top view (third row). Colors code is as follow: Mutations (Red), S1 subunit (Cyan), S2 subunit (Green), Furin Cleavage Site (Yellow), Fusion peptides FP1 and FP2 (Black), and the arrows show the TMPRSS2 cleavage site (Orange).

### 3.4. Effect of omicron S protein mutations on the immunogenicity

Non-synonymous mutations of the different SARS-CoV-2 variants caused changes on the epitope probability and antibody exposure hence immunogenicity. Most of the epitope changes are noticeable in the S1 domain of the spike protein. Figure 4 shows the percentage of the exposed epitope on the different variants. Detailed probability and exposure states of each residue as predicted by BepiPred are listed in Supplementary Table 1.

**Figure 4.**
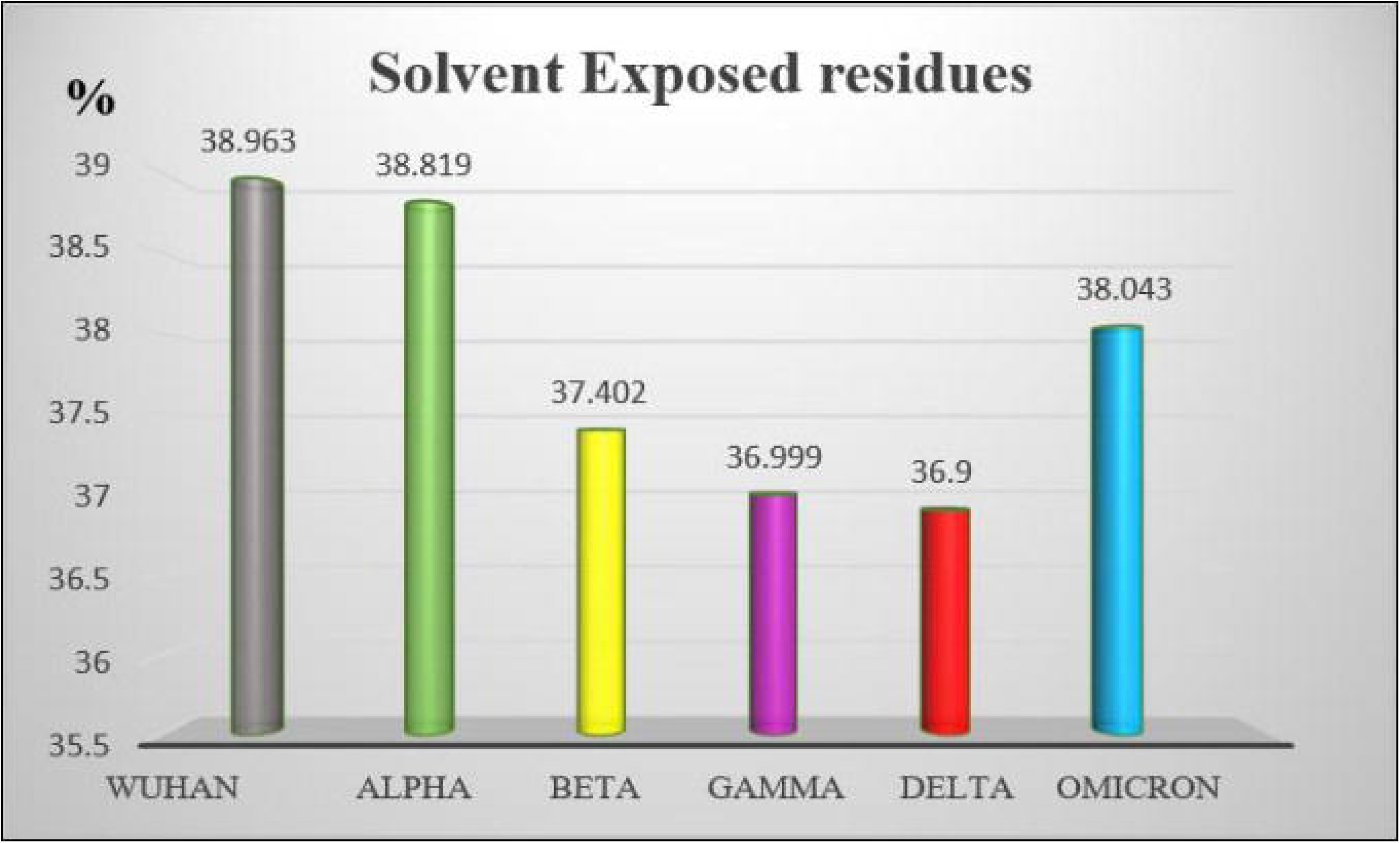
The percentage of solvent exposed residues in the S protein of the five SARS-CoV-2 VOCs.

### 3.5. Effect of omicron S protein mutations on the 3D structure of the molecule: Loop modeling and quality assessment

Computational methods allow the building of accurate protein models of the SARS-CoV-2 S protein based on data input and the alignment with experimentally solved multiple templates molecules. We used the Phyre2 web server to generate 3D models for the extracellular domain of the variants α, β,γ,δ and O monomers spike protein (Fig 5). The structures were generated with 100% confidence and 84% coverage for *alpha, beta and omicron*, and 83% coverage for *gamma* and *delta* by the single highest scoring template. The quality assessment of these models obtained by superimposition with the crystalized structure of the SARS-CoV-2 spike glycoprotein (closed state) 6VXX Chain B, showed an RMSD of 1.086 for *alpha*, 1.085 for *beta*, 1.079 for *gamma*, 1.075 for *delta* and 1.075 for *omicron* (Fig. 5). The Tm-scores were all less than 1 and both RMSD and TM-scoring values are acceptable and indicates high similarity between the crystalized model and the generated one showing the same folds. In addition, CMView common contact map gave a high similarity score ranging between 80% - 81.5% keeping in mind that the crystalized structure is missing few residues (gaps) including the furin cleavage loop. What is important here is that these generated models include the furin cleavage site that is lacked in all the crystalized models deposited on the protein database (PDB). Moreover, comparison of the contact map percentage and TM-align scoring gave an indication of the similarity between the different models (Table 4). Given the highly reliable Tm score and RMSD values, the computationally generated models are suitable to be used for further analysis especially for evaluating the effect of mutations on and around the furin cleavage site.

**Figure 5.**
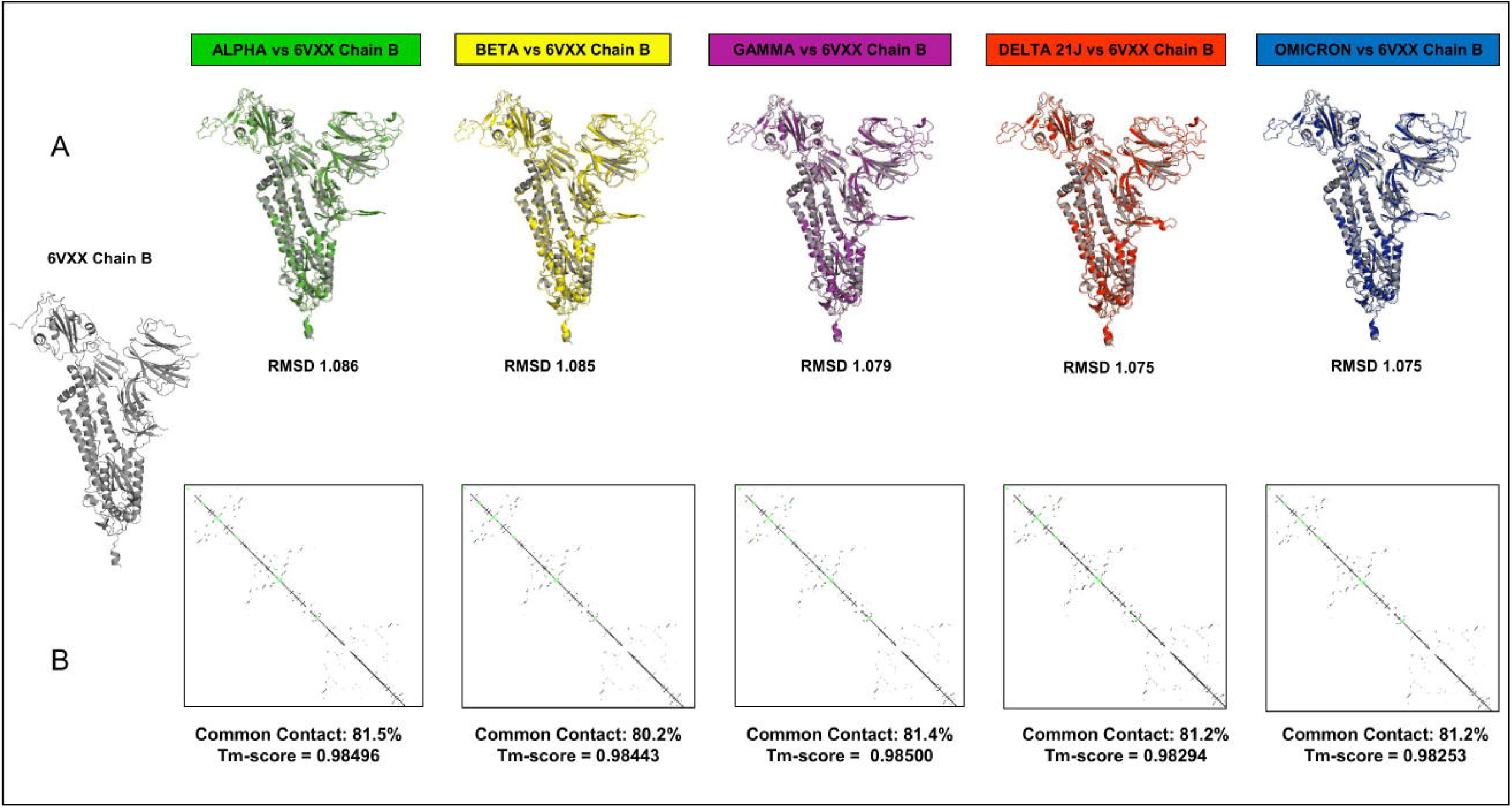
This figure represents the 3D models generated for the α, β,γ,δ and O variants, their structural analysis and the quality assessment by superimposition with the chain B of the crystalized model PBD 6VXX (A) Superimposition RMSD values (B) Calculated common contact percentages and Tm-scores.

**Table 4.**
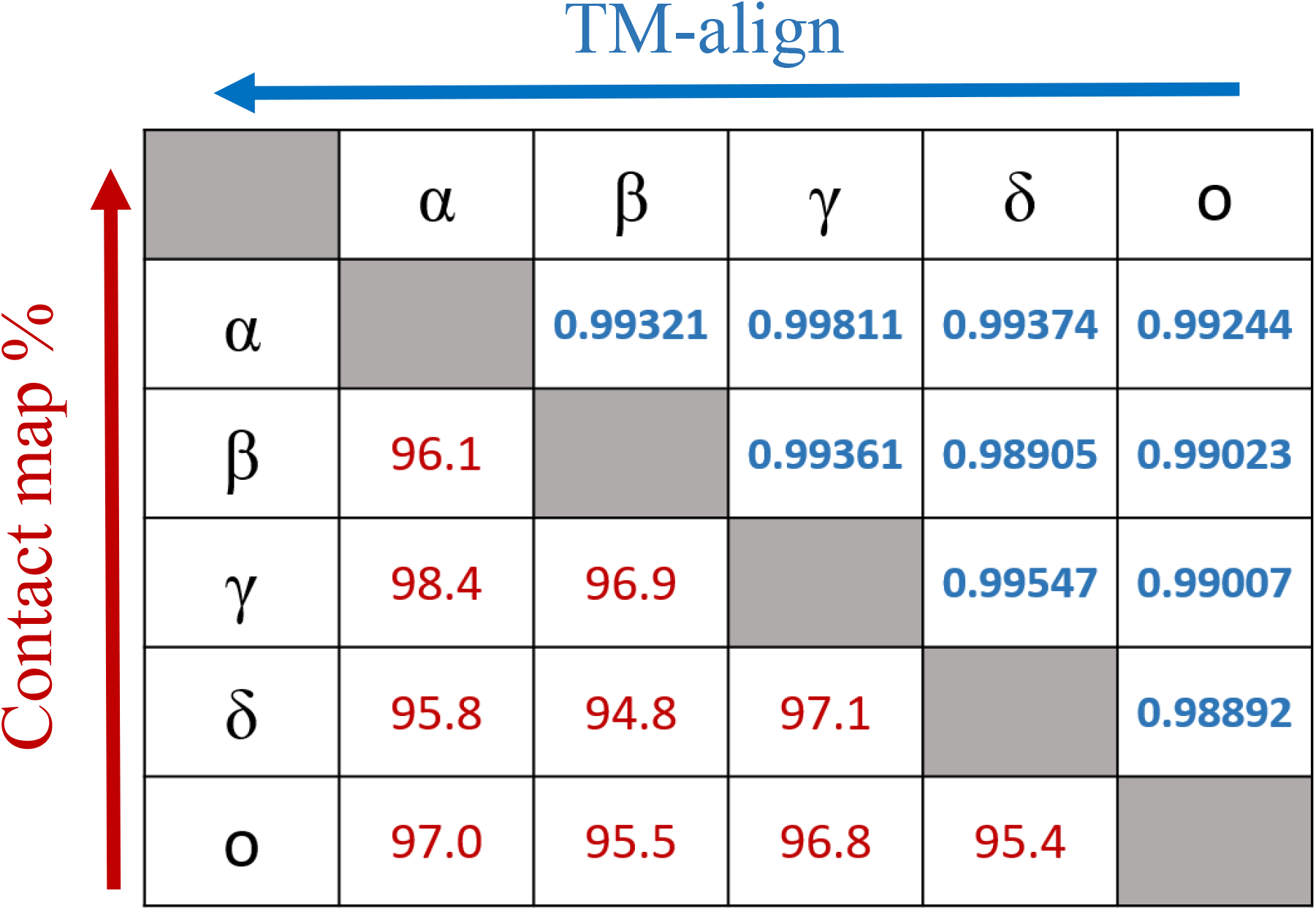
Comparison of contact map percentage (Red) and TM-align (Blue) scoring between the different SARS CoV-2 VOCs. The arrowheads denote the variant used as a reference when calculating the data.

### 3.6. The effect of contact residues’ mutations on the SARS-CoV-2 S protein/ACE2 complex thermodynamic stability

Sequence analysis showed that *omicron* variant has the most mutated RBD with 15 different mutations out of which, nine mutations were in the contact residues with ACE2 receptor. The *alpha* variant showed only one mutation in the contact residues (N501Y), *beta* and *gamma* showed three mutations all are of the contact residues while *delta* showed two mutations none of which are in the contact residue with ACE2. Analyzing theses mutations as single mutation showed different effect on the complex of SARS-CoV2 RBD with ACE2 receptor. Where some mutations showed to be deleterious as single mutation, others have stabilizing effect. However, the combination of several mutations in the contact residue shows different effect; in the case of the *omicron* variant, the combination of 9 mutations in the contact residues showed to be not deleterious even though they slightly destabilize the complex with ACE2 with an accumulative ΔΔG= 0.18. *beta* and *gamma* variant with three mutations in the contact residue showed an accumulative ΔΔG of 1.79 and 1.07 respectively, which highly destabilizes the complex and decrease the binding affinity. *Delta* has only two mutations that occurs out of the contact residues with a stabilizing effect on the complex and hence increasing affinity to ACE2 and an accumulative ΔΔG= −0.33. The single contact residue mutation of the *alpha* variant (N501Y) showed destabilizing effect that we reported previously (Ashoor et al., 2021) (Table 5).

**Table 5.** Comparison of the mutations profile of five SARS-Cov-2 VOCs S protein in the contact residues with the ACE2 receptor. Positive and negative signs correspond respectively to destabilizing (decreases in binding affinity) and stabilizing mutations (increases in binding affinity).

| 6LZG<br>Contact<br>residues | *Alpha*<br>B.1.1.7 | *Beta*<br>B.1.351 | *Gamma*<br>P.1 | *Delta*<br>B.1.617.2 | *Omicron*<br>BA.1 | Single<br>mutations',<br>$\Delta\Delta G$ | Effect of<br>single<br>mutation |
| --- | --- | --- | --- | --- | --- | --- | --- |
| K417 |  | K417N |  |  | K417N | 0.75 | Destabilizing |
|  |  |  | K417T |  |  | 0.18 | Destabilizing |
| G446 |  |  |  |  | G446S | -0.18 | Stabilizing |
| G447 |  |  |  |  |  |  |  |
| Y449 |  |  |  |  |  |  |  |
| - |  |  |  | L452R |  | -0.57 | Stabilizing |
| Y453 |  |  |  |  |  |  |  |
| L455 |  |  |  |  |  |  |  |
| F456 |  |  |  |  |  |  |  |
| Y473 |  |  |  |  |  |  |  |
| A475 |  |  |  |  |  |  |  |
| G476 |  |  |  |  |  |  |  |
| S477 |  |  |  |  | S477N | -0.15 | Stabilizing |
| - |  |  |  | T478K |  | 0.19 | Destabilizing |
| E484 |  | E484K | E484K |  | E484A | 0.17 | Destabilizing |
| G485 |  |  |  |  |  |  |  |
| F486 |  |  |  |  |  |  |  |
| N487 |  |  |  |  |  |  |  |
| Y489 |  |  |  |  |  |  |  |
| F490 |  |  |  |  |  |  |  |
| L492 |  |  |  |  |  |  |  |
| Q493 |  |  |  |  | Q493R | 0.25 | Destabilizing |
| Y495 |  |  |  |  |  |  |  |
| G496 |  |  |  |  | G496S | 1.2 | Destabilizing |
| F497 |  |  |  |  |  |  |  |
| Q498 |  |  |  |  | Q498R | 2.34 | Destabilizing |
| P499 |  |  |  |  |  |  |  |
| T500 |  |  |  |  |  |  |  |
| N501 | N501Y | N501Y | N501Y |  | N501Y | 1.53 | Destabilizing |
| G502 |  |  |  |  |  |  |  |
| V503 |  |  |  |  |  |  |  |
| Y505 |  |  |  |  | Y505H | 1.33 | Destabilizing |
| Q506 |  |  |  |  |  |  |  |
| **Accumulative**<br>**$\Delta\Delta G$** | 1.53 | 1.79 | 1.07 | -0.33 | 0.18 |  |  |
| **Effect of**<br>**accumulative**<br>**mutation** | Destabilizing | Destabilizing | Destabilizing | Stabilizing | Destabilizing |  |  |

### 3.7. Effect of mutations in the S protein of SARS-CoV-2 major variants on the polar interactions with the ACE2 receptor

Using the crystalized structure PDB ID 6LZG that represent the interaction between SARS-CoV-2 receptor binding domain with ACE2 receptor as a reference, we generated models for the five variants on the LigPlot+ software and analyzed the polar interaction patterns. The interacting residues and type of interactions are listed in Table 6. All the variants have one or more mutations on the contact residues except for the *delta* variant. At a first glance to Table 6 and by comparison, one can spot the interaction pattern similarity between *alpha* and *delta* variants despite the fact that the *alpha* variant showed only one mutation in the contact residue (N501Y) and *delta* has two mutations out of the contact residues (L452R and T478K). Both *alpha* and *delta* variants were reported to be highly transmissible with *delta* being 60% higher (Duong, 2021). *Delta* shows that eight out of 9 main polar interactions are present with one missing polar interaction (Spike/ACE2: Asn487/Tyr83) and the addition of new two polar interactions (Gln493/Glu35 and Tyr505/Glu37) and one salt bridge (Glu484/Gln24). Similarly, *alpha* is missing the same polar interactions but shows the same new salt bridge Glu484/Lys31 and no additional new polar interactions. The addition of new polar interactions and the Glu484/Lys31 salt bridge in the *d*elta variant could be the cause of the stabilizing effect on the complex with ACE2, which can explain the high transmissibility.

**Table 6.**
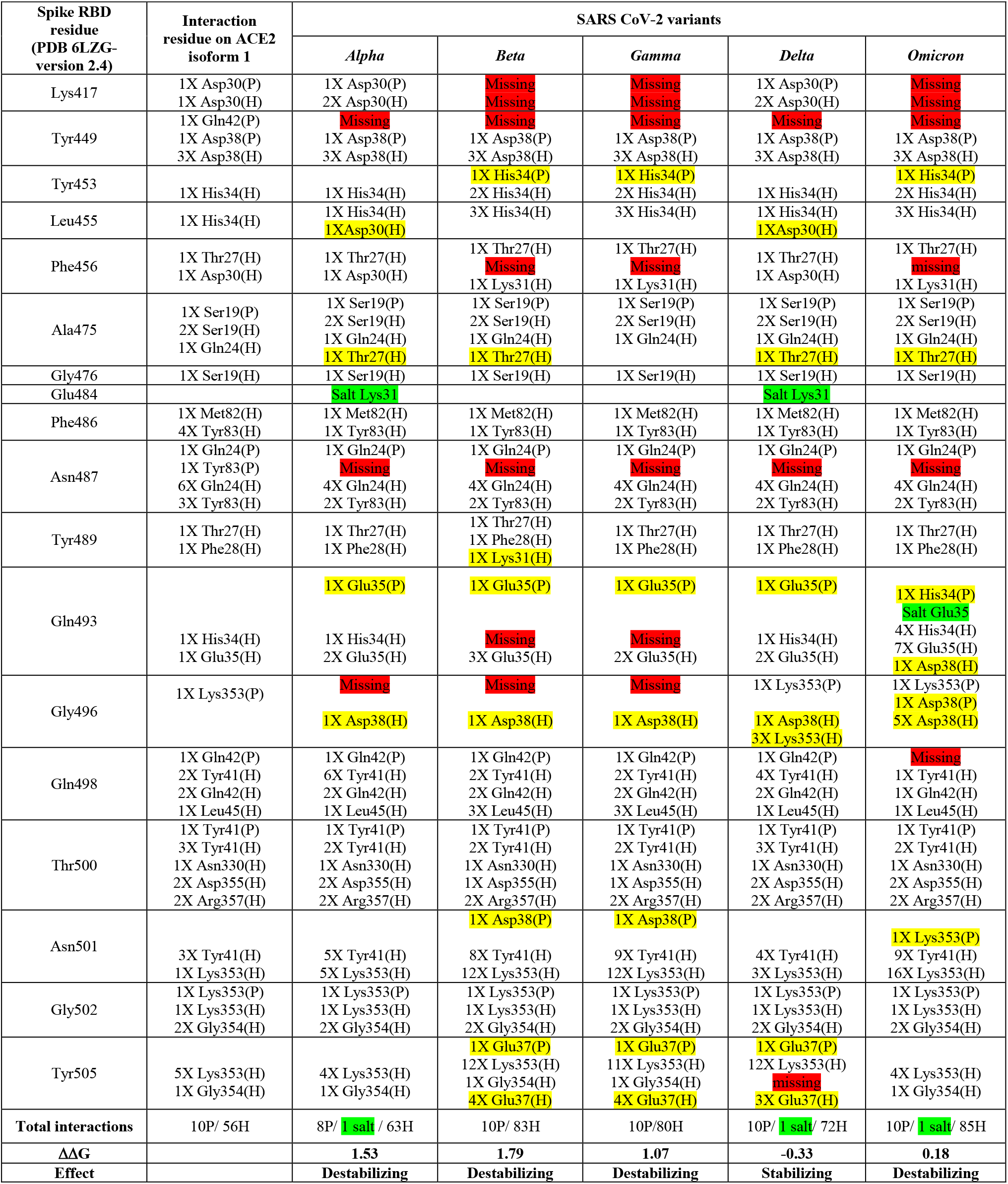
Contact residues’ interaction pattern of the S protein/ ACE2 complex of different SARS CoV-2 variants. Polar interactions (P), hydrophobic interactions (H), Missing interactions are highlighted in red, new interactions highlighted in yellow, salt bridges are highlighted in green.

In addition to Glu484, other mutations on RBD binding hot spots residues, have been also linked to antibody binding and neutralization including mutations on Lys417, Gly446, Phe456, Asn501, Gly477 and Asp614 (Greaney et al., 2021a; Liu et al., 2021b; Montefiori, 2021; Supasa et al., 2021). Interestingly, *omicron* variant compiles all these mutations (Table 5) indicating a potential higher degree of immune escape. Additionally, *omicron* has a consecutive mutations on residue Gln493, Gly496 and Gln498 that dramatically affect the interaction with ACE2. These residues are structurally in the receptor binding ridge and Gln493 is known to form with Leu455 the two receptor binding motif (RBM) stabilizing hot spots and to be a target for some therapeutic antibodies (Shang et al., 2020; Greaney et al., 2021b). This interaction disrupted profile may sum up into less stable complex with ACE2 (ΔΔG= 0.18) (Table 6) and loss of antibody neutralization. Furthermore, the comparison of the interactions’ differences between SARS-CoV-2 variants showed some interactions that are conserved in all the variants suggesting their importance in the complex stability.

## 4. DISCUSSION

Despite significant global health care measures and social mitigation efforts along with the availability of a number of vaccines, the SARS-CoV-2, COVID-19 pandemic entered its 3^rd^ year and the numbers of cases are soaring worldwide (https://www.gisaid.org/, https://coronavirus.jhu.edu/map.html). This is mainly due to the emergence of new and more fitted viral variants that keep fueling this pandemic. To create awareness about any additional health issue a new variant may cause, WHO the world most influential health agency, has developed a labeling system to classify the new variants, in addition to the conventional nomenclature. The CDC has adopted this labeling system but introduced major changes. Among the labels used by WHO and the CDC is the label “Variant Of Concern or VOC”. The word “concern” is synonymous of anguish, anxiety and apprehension. It naturally enjoins fear and disturbance, which predispose to take immediate actions and make odd changes. Indeed, when a new variant is labeled VOC, countries are inclined to close their borders and take drastic counter measures. This was particularly striking when a number of countries banned travels from South Africa that was the first country to describe and report the genomic sequence of the *omicron* variant. While preventive and cautionary actions are mandatory to control pandemics, these should be based on solid scientific evidences. However, it takes time and coordinated efforts for the scientific community to generate the data needed to label accurately new variants according to criteria such as transmissibility, disease severity, and change in the epidemiological pattern, immune escape, and resistance to previously neutralizing antibodies, efficacy of existing therapies or vaccine efficiency. Meanwhile, the rapid pandemic progress requires timely response including a good communication system. In this study, we highlight some flaws in the labeling system developed by WHO mainly to ease communication with the national health authorities and the public. Indeed, in addition to the non-appropriate wording for labeling emergent variant, the combination of criteria used to define a variant label as shown in WHO and the CDC web sites, is not accurate. Indeed, in their definitions of the different labels, both agencies build a combination of criteria using the prefix “OR” but not “AND”. This introduces a confusion that is amplified by the CDC use of more labels, and different formulation and different combinations of the criteria used to attribute a label to a new variant. According to WHO a label is supposedly assigned to a variant through a comparative assessment with the previous ones. Even though SARS-CoV-2 variants are primarily detected upon the virus genomic sequence changes and particularly mutations in important functional regions, the genetic variations do not clearly appear among the viral attributes WHO and the CDC use to formulate their definition of the different variants. The labeling system developed by WHO and the definitions of the criteria for each label used by both WHO and the CDC do not mention a comparative assessment of the genetic variations but rather retain clinical, epidemiological, transmissibility and other non-genetic viral attributes. Such parameters need a substantial time to be accurately determined. It seems that the current labeling system relies more on fear from emergent genetic variations “the super killer virus” than evidences about the impacts of mutations (Caini et al., 2018). Thus, it seems premature and not appropriate to give a new variant the VOC label. Even when a greater transmissibility can be established, this does not necessarily mean greater severity. For example, the very transmissible H1N1 influenza virus variant was not as severe as many other influenza viruses and the epidemic has naturally faded away (Grubaugh et al., 2020). With reference to WHO variant labeling system H1N1 could have been labeled VOC while it has never been one. Therefore, to solve the dilemma between the necessity of rapidly labeling new variant and the lack of scientific evidences, a good way is to analyze thoroughly the genetic variations observed to generate rapidly the best data on the potential attributes of the new viral variant. To achieve this analysis, the computational prediction of variant effect on protein stability, function and interaction is a very useful way to determine the variant “importance”. Several methods can be used based on the availability or not of 3D structure of the protein. In this study, we show that computational prediction and *in silico* comparative studies between new variants and older more characterized ones, can provide good insights into the impact of the mutations and thus the potential behavior of the emergent variants. Indeed, the observation that most of the mutations in SARS-CoV-2 VOCs map to the solvent exposed regions along with the comparison of antigenicity predictions, represent a good basis to make a fair assumption on a potential immune escape and/or a likely reduction of antibodies neutralization and to anticipate an evaluation of vaccine effectiveness. In addition, the comparative analysis of the S protein 3D structure between the five SARS-CoV-2 VOCs showed an identical overall folding as demonstrated by the RMSD values and Tm-scores of each model. Meanwhile the comparison of contact residues maps gave a fine tuning of the structural divergence which suggest potential functional differences that need to be further analyzed.

Furthermore, computational studies of the mutations effect on the thermodynamic stability of the S protein /ACE2 complex and the comparative analysis of the pattern of polar interactions of the different variant with the ACE2 receptor, give good indications to predict transmissibility and virulence and draw some plausible epidemiological scenario. Indeed, the effect of mutations can be computed as single or combined mutations. The data we obtained with the combined mutations of the *omicron* variant that has a much higher number of mutations than the other variants, shows that this variant engages into a more stable interaction with ACE2 than the β and γ variants do and has more interactions than the delta variant.

For the other variant labeling criteria, several studies discussed the importance of the Glu484Lys mutation in the interaction with ACE2 and immune escape (Greaney et al., 2021b; Makdasi et al., 2021). This was also observed with the *beta* and *gamma* variant that carry Glu484Lys mutation. It was reported that the *beta* variant have reduced antibody neutralization compared to the *delta* variant. Besides *beta’s* resistance to neutralizing antibodies increased by 9.4 fold to convalescent plasma and 10.3 to 12.4 fold for sera from individuals who have been vaccinated (Liu et al., 2021a). In addition, it was suggested that new variants with the same mutation might bear new challenges for current vaccines or monoclonal antibody therapies (Krause et al., 2021; Zhou and Wang, 2021).

Meanwhile our analysis of the pattern of polar interactions between the different variants shows that *omicron* has an extra 13 hydrogen bonds as compared to variant *delta*. In addition, we noticed the presence in the more transmissible *alpha* and *delta* variants of an extra salt bridge between the S protein Glu 484 and ACE2 lysine 31 and the presence of an extra salt bridge between the S protein Gln 493 and ACE2 Glu 35 in the *omicron* variant. This observation combined to the observed high number of H bonds and the thermodynamic stability data, predicts a potential more efficient entry into host cells and enhanced transmissibility of the *omicron* variant, which has been confirmed since the description of *omicron* mutations profile. Nevertheless, relying on the study of a single mutation or the computing of one biophysical feature of a variant structure to predict how a viral attribute would evolve is not sufficient. Furthermore, multiple genes can control epidemiologically relevant viral attributes such as the mode of transmission and virulence. Therefore, it is recommended to integrate the data on multiple mutations with computing of various viral structural features to make the best predictions and attribute the right label to a given variant.

In conclusion, the system of SARS-CoV-2 labeling developed by WHO and amended by the CDC has some major flaws. Relying on the integrated biophysical and structural data generated from computational comparative predictions of the likely behavior of a new variant would help in the rapid and accurate labeling of emergent variants. Meanwhile, given our provisional and incomplete knowledge and the uncertain nature of the COVID-19 pandemic, it would be wise to operate in epistemic self-abnegation, use the best tools and knowledge we have at hand and introduce revisions whenever new evidences become available.

## Author Contributions

Dana Ashoor: *In Silico* analysis, methodology, data curation, writing and editing, Maryam Marzouq: Mutations’ review, illustrations figures and tables, revision. Khaled Trabelsi: Data retrieval from data banks and formatting, Sadok Chlif: Analysis and illustration of the WHO and CDC variants-labelling criteria, Nasser Abotaleb: Data retrieval and iconography, Noureddine Ben Khalaf: Revision of the *in silico* study, Ahmed R. Ramadan: Data cross checking and organization. M-Dahmani Fathallah: Project conception, work design, data analysis, writing, editing, and supervision

